# *ZC4H2* loss of function is associated with temporal dysregulation of neural stem proliferation and neuron development

**DOI:** 10.64898/2026.01.15.699794

**Authors:** Yomira Palacios, Camille Loret, Viktoria Haghani, Kyle D. Fink, Julian A.N.M. Halmai

## Abstract

*ZC4H2* is an X-linked zinc finger transcription factor essential for early neurodevelopment. Pathogenic variants in *ZC4H2* are associated with both central and peripheral nervous system pathologies. The molecular and cellular mechanisms driving these phenotypes remain poorly understood, particularly in human female models that have undergone X chromosome inactivation. Neuronal models were differentiated from a human female cell line with a de novo Xq11.2 deletion causing *ZC4H2* loss of function, associated with arthrogryposis multiplex congenita and cognitive impairment. Using iPSC-derived neural stem cells and cortical organoids, we identified premature neuronal differentiation, reduced BMP-SMAD signaling, and decreased SMAD1/5 phosphorylation. In cortical organoids, *ZC4H2* deficiency altered neurogenesis timing, retaining proliferative progenitors while prematurely activating neuronal programs, leading to enlarged organoids with persistent dysregulation of gene programs required for complete neuronal maturation. We identified *ZC4H2* target genes likely to mediate these phenotypes and tested a codon-optimized transgene showing both the restoration of SMAD1/5 phosphorylation, *BMPR2* gene expression, and improved neuronal complexity. These results demonstrate effective *ZC4H2* restoration in complex human models and highlight therapeutic potential for *ZC4H2*-linked neurodevelopmental disorders.

## Introduction

*ZC4H2*-associated rare disorders (ZARD) are ultra-rare X-linked neurodevelopmental disorders caused by pathogenic variants in the *ZC4H2* gene, which encodes a zinc finger transcription factor critical for both central and peripheral nervous system development. Clinically ZARD is frequently associated with arthrogryposis multiplex congenita (AMC), seizures, intellectual disability (ID), and brain malformations such as microcephaly and macrocephaly(1,2). Although few patient cohort studies exist, a recent study of 40 individuals with confirmed *ZC4H2* variants revealed significant genotype-phenotype and sex differences (3). Females with *de novo* loss-of-function (LoF) variants commonly exhibit profound cognitive impairment and musculoskeletal deficits, while males with maternally inherited missense variants showed a higher prevalence of seizure phenotypes. Although studies have shown significant differences in symptom severity between males and females, certain mutations like *de novo* deletion mutations within females result in more serve disease manifestations including cognitive impairment. The exact roles these genetic variants play in contributing to cognitive impairment, especially within the female ZARD population remains unknown and can offer insight into potential corrective strategies for therapeutic intervention.

Mouse studies have determined that *Zc4h2* is highly expressed during early neurodevelopment and declines postnatally, supporting that *ZC4H2* plays an important role in early brain development (4). Functional studies have shown that *ZC4H2* localizes to the postsynaptic density of excitatory neurons were overexpression of altered *ZC4H2* reduces dendritic spine density and number, while other studies looking at conditional knockout of *ZC4H2* in forebrain excitatory neurons increases neuronal complexity resulting in impaired cognitive functions including learning and memory(5,6). Regarding GABAergic neuron development, zebrafish knockout models exhibit hyperactive behavioral phenotypes accompanied by a reduction in GABAergic interneuron populations (7). Together, these findings suggest that *ZC4H2* plays critical roles in the development of both excitatory and inhibitory neurons. Transcriptomic analysis of differentially expressed genes (DEGs) in knockout mouse Neural Stem Cells (NSCs) indicates that *Zc4h2* regulates pathways involved in synapse assembly, telencephalon development, neural precursor proliferation, and axon development(8). Knockdown and knockout studies further demonstrate that *ZC4H2* is essential for NSCs survival and proliferation(5,8). Notably, *Zc4h2* deficiency is associated with decreased NSCs proliferation and increased differentiation, indicating a shift in the normal balance between NSC maintenance and neurogenesis, supporting dysregulated neurodevelopment. Mechanistic studies in *Xenopus* have found that *ZC4H2* acts as a cofactor for other transcription factors. Specifically, *ZC4H2* stabilizes Smad1 and Smad5 by preventing their ubiquitination and degradation by Smurf1 and Smurf2 (9). The BMP-SMAD signaling cascade is essential for regulating GABAergic neuron specification, progenitor self-renewal, and neuronal differentiation (7,9–11). Dysregulation of this pathway has independently been shown to accelerate neural stem cell differentiation and to disrupt GABAergic interneuron development, leading to network hyperactivity (15,17). Since *ZC4H2* deficiency and Smad1/5 dysregulation share features such as network hyperexcitability, increased neuronal differentiation, and impaired GABAergic interneuron development, investigating the link between *ZC4H2* and Smad1/5 in ZARD is critical. Despite these insights, the role of *ZC4H2* in human female patient-derived neural models remains unexplored.

In this study, we generated and characterized NSCs and cortical organoids (CtOs) from induced pluripotent stem cells (iPSCs) derived from a female patient carrying a heterozygous 244-kb microdeletion encompassing the transcription start site, promoter, and exon 1 of *ZC4H2*. This pathogenic variant, clinically associated with AMC, seizures, and mild cognitive impairment, resulted in accelerated NSCs proliferation and the formation of significantly larger CtOs compared to controls. Transcriptomic profiling of both NSCs and CtOs revealed temporal dysregulation of stem-cell maintenance and neuronal differentiation programs, indicative of premature neuronal development coupled with dysregulated maturation. Knockdown of *ZC4H2* in wild-type NSCs confirmed *BMPR2* and *MAP2* as *ZC4H2* targets, whereas transgene restoration of *ZC4H2* rescued BMPR2 expression and SMAD1/5 phosphorylation, mechanistically establishing that *ZC4H2* regulates BMP-SMAD signaling cascade activation. Morphological analyses demonstrated that *ZC4H2* is required for maintaining neuronal structural complexity and regulating NSCs proliferation, which was improved upon gene restoration. Collectively, these findings define novel and reversible molecular and cellular phenotypes associated with *ZC4H2* deficiency, identify *MAP2* and *BMPR2* as potential therapeutic targets, and highlight the BMP-SMAD signaling cascade as a key pathway disrupted in ZARD.

## Results

### ZARD NSCs exhibit accelerated proliferation and increased cell cycle activity

To investigate the pathophysiological consequences of a 244kb microdeletion (ARR[HG19] XQ11.2(64,170, 839-64,414573) X1) in *ZC4H2* **(Figure 1A),** we generated NSCs and CtOs from a patient-derived iPSC line (GM28603 (**Figure 1B**). RT-qPCR analysis revealed a 2.8-fold reduction in *ZC4H2* expression in ZARD NSCs compared to IMR90 sex-matched control NSCs (*p* < 0.05) (**Figure 1C**), confirming LoF in patient-derived cells. Publicly available datasets indicate that *ZC4H2* does not escape X chromosome inactivation, supporting that patient derived NSCs are mosaic. Previous reports in mouse models have shown that KO of *Zc4h2* reduces NSCs proliferation, whether this phenotype is associated with this variant remains unknown(5,8). To investigate the differences in growth kinetics between variant and control NSCs, we conducted a series of proliferative assays. Growth curve analysis revealed that ZARD NSCs proliferate more rapidly than sex-matched control IMR90F (**Supplement Figure A**). ZARD NSCs exhibited a significant increase in EdU+ DNA incorporation compared to both IMR90F and 80230F NSCs control lines, corresponding to a 50% and modest 3% increase, respectively demonstrating an increase in NSCs proliferation (**Figure 1D**). Additional studies assessing cell cycle entry via Ki-67 immunocytochemistry showed a ∼20% increase in KI-67^+^ cell in ZARD NSCs versus control IMR90 NSCs (P < 0.05), indicating an increased cell cycle activity. Notably, Ki-67 expression in ZARD NSCs was not significantly different from that observed in 80230F cells (**Figure 1E, F**). These results demonstrate that this variant is associated with accelerated NSCs proliferation. To examine differentially expressed genes (DEGs) and pathways with variant NSCs, that could further explain this, we performed RNA sequencing analysis. Differentially expressed genes were identified using limma-voom with Benjamini-Hochberg correction, applying a significance threshold of FDR < 0.05 without an additional fold-change cutoff. ZARD variant NSCs exhibited widespread transcriptional changes, with 6038 downregulated and 6365 upregulated genes relative to IMR90F NSCs. The top five upregulated genes by fold change were *ZIC3* (log2 9.74), *CACNG7* (log2 9.75), *FABP7* (log2 9.78), *IRX2* (log2 9.96), and *CEACAM21* (log2 9.91). Among these, the neurodevelopmental regulators *IRX2*, *FABP7*, *CACNG7*, and *ZIC3* showed consistent dysregulation, implicating disruptions across key developmental signaling pathways FGF-ERK and WNT (*IRX2*), BMP/TGF-β (*ZIC3*), and neuronal maturation and calcium channel regulation (*CACNG7*, *FABP7*). In contrast, CEACAM21 has no well-established role in neurodevelopment. Among the most strongly downregulated genes, *INHBA* (log2 -10.56) is a key TGF-β/Activin regulator involved in neural progenitor maintenance and differentiation, whereas *IL7R* (log2 -11.02) is an immune receptor with no known neurodevelopmental function. Other markedly reduced genes *MEDAG* (log2 -10.65) and *KRT7* (log2 -10.28) lack neuronal relevance, while SHISA3 (log2 -9.82) participates in WNT/FZD modulation during telencephalic patterning **(Supplemental Figure B).** The top 50 upregulated genes with a log FC of 2 and a *p*<0.05 were filtered into GO analysis. GO-term biological processes revealed ZARD NSCs exhibited a significant upregulation in pathways involving stem cell development and stem cell differentiation (**Figure 1G**). When looking at DEGs important for NSCs proliferation and self-renewal, *MSI2* (log2 2.56), *TRIM71* (log2 5.08), and *SOX2* (log2 9.02) were enriched consistent with a hyperproliferative state **(Supplemental Table 2)**. Together, these results demonstrate that this ZARD variant exhibits accelerated NSCs proliferation accompanied by dysregulated genes important for stem cell development and self-renewal, suggesting abnormal expansion of the NSCs in ZARD.

**Figure 1:**
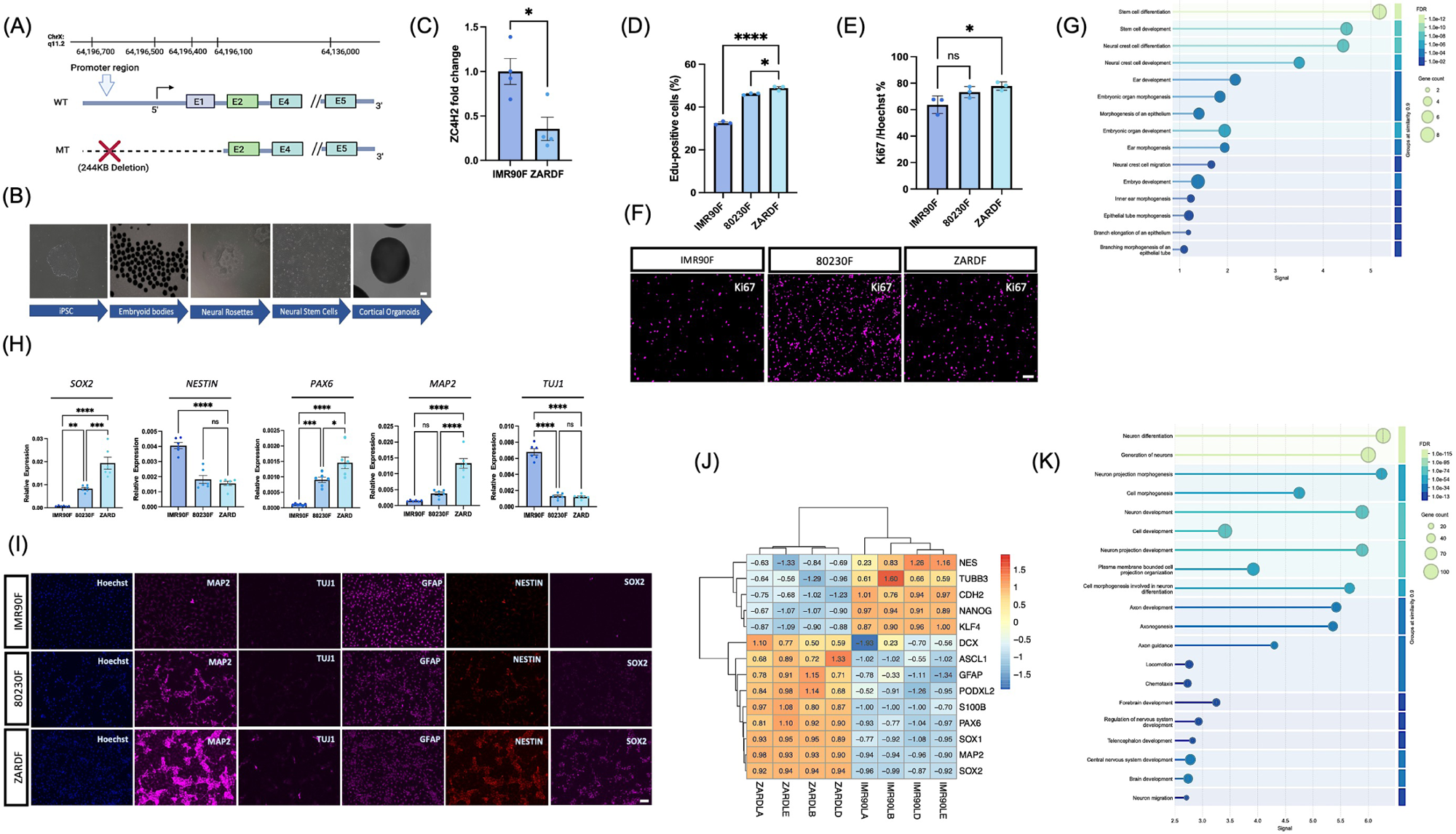
Microdeletion in *ZC4H2* promoter affects gene regulation and neural phenotypes in ZARD models. (A) Schematic illustrating the microdeletion spanning the ZC4H2 promoter region in the model system, which prevents sgRNA binding and transcriptional activation of the mutant allele. (B) Representative image demonstrating successful differentiation of patient-derived iPSCs into NSCs and CtOs. (C) RT-qPCR analysis showing significant downregulation of *ZC4H2* gene expression in variant NSCs compared to sex-matched control IMR90F NSCs. unpaired t-test (p value 0.0167). (D) EdU incorporation assays demonstrate a higher proportion of proliferating cells in ZARD NSCs (n=3). Percent EdU⁺ cells were determined by flow cytometry as the proportion of events within the live singlet gate falling above the EdU fluorescence threshold defined using unstained (EdU⁻) controls. (E-F) Ki67 (magenta) ICC of NSCs from IMR90F, 80230F, and ZARDF lines and subsequent quantification of of protein intensity in NSCs reveals elevated Ki67 staining in ZARD NSCs compared to IMR90 control. % Ki67⁺ cells (mean ± SEM, n=3 biological replicates. Statistical analysis was performed using one-way ANOVA with Dunnett’s multiple comparisons test. ZARDF vs. IMR90F, *p* value 0.0217; ZARDF vs. 80230F, not significant (*p* value 0.4595). Scale bar, 100 μm. (G) Dot plot showing GO Biological Process enrichment for upregulated pathways associated with the top 50 upregulated genes (log2 fold change ≥ 2, adjusted *p* <0.05). The x-axis represents enrichment significance (-log10 FDR). Dot size indicates the number of genes associated with each GO term, and dot color represents FDR. Enriched terms represent biological processes overrepresented among upregulated genes. (H) Relative mRNA expression analysis showing increased levels of neural stem cell (NSC) and neuronal markers in mutant ZARD NSCs compared to sex-matched controls (IMR90 and 80230F). (I) Immunocytochemistry (ICC) demonstrating differential protein expression of select NSC and neuronal markers between mutant and sex-matched control NSCs. (J) Heat map showing row-scaled (z-score) log-CPM expression values of genes involved in iPSC, NSC, and neuronal development across control and ZARD NSCs. Red and blue indicate higher and lower expression relative to each gene mean, respectively. Hierarchical clustering was performed on genes and samples. (K) GO Biological Process enrichment dot plot highlighting the upregulated pathways involved in neurogenesis and neuronal differentiation within ZARD NSCs corresponding to the top 500 upregulated genes (log2 fold change ≥ 2, adjusted *p* <0.05 in ZARD NSCs. Dot size corresponds to gene count, dot color to FDR (high to low). The x-axis shows -log10(FDR). GO terms are grouped by semantic similarity to highlight related biological processes. All mRNA transfection experiments have between 3-6 replicates. Significance testing for quantification analyses conducted by both unpaired t-test and one way ANOVA with statistical significance indicated as \*\*\**p* < 0.0001, \*\**p*< 0.001 *p < 0.05.

### ZARD NSCs display transcriptional and phenotypic signatures of a proneuronal fate

To further delineate whether ZARD NSCs are prematurely differentiating into neurons, we examined differences in the expression of NSCs and neuronal markers on a transcript and protein level. Interestingly, ZARD NSCs show an increase in the expression of immature *ASCL* (log2 3.98) and mature neuronal markers including *POU3F2* (log2 8.31), *GPM6A* (log2 4.12), *SNCA* (log2 2.45), and *MAP2* (log2 2.75) on an mRNA level **(Supplemental Table 2).** To validate the transcriptomic dysregulation of neuronal and NSC markers a RT-qPCR panel was conducted within ZARD NSCs compared to sex-matched controls IMR90 and 80230F. Mature neuronal marker *MAP2* and neural stem cell markers *SOX2* and *PAX6* were significantly upregulated in ZARD NSCs compared to IMR90 and 80230F controls (p < 0.05) (Figure 1H). When compared to IMR90 controls, the fold changes for *MAP2*, *SOX2*, and *PAX6* were 9.18, 26.46, and 14.53 respectively. In comparison to 80230F controls, the fold changes were lower but still indicated significant upregulation in ZARD NSCs, with values of ∼3.52, 2.34, and 1.61 for *MAP2*, *SOX2*, and *PAX6* on average, respectively. This confirms that this dysregulation was consistent when comparing variant NSCs across multiple controls. Notably, when examining protein expression levels via ICC, we observed a trend toward increased neuronal marker MAP2 expression in ZARD NSCs compared with IMR90F controls ∼2.1-fold on average, although this difference did not reach statistical significance (**Supplemental Figure C**). This pattern suggests a potential increase in neuronal maturation in ZARD NSCs relative to controls (Figure 1I). Furthermore, Gene Ontology (GO) analysis revealed enrichment for biological processes associated with neuron differentiation (FDR 1.0E-115), generation of neurons (FDR 1.0E-115), and Neuron development (FDR 1.0e-74), further supporting a premature neuronal fate in these rapidly proliferating NSCs (**Figure 1K**). To examine the potential shift in cell fate we examined the changes in gene expression of astrocytic markers in ZARD NSCs. Bulk RNA-sequencing data and ICC analysis revealed an increase in expression of S100B and GFAP at both the mRNA and protein levels, indicating a potential shift towards astroglial lineage commitment (**Figure 1I, J).** Together, these findings confirm that ZARD NSCs exhibit both premature neuronal differentiation and increased astrocytic marker expression, further supporting their altered developmental trajectory.

### *ZC4H2* LoF results in the dysregulation of the SMAD-BMP signaling cascade

The BMP-SMAD signaling cascade is essential for regulating cell fate specification and maintaining the balance between NSCs self-renewal and differentiation. Previous studies have described *ZC4H2* to play an important role in stabilizing SMAD1/5 from degradation. Here we aimed to understand the consequences of SMAD1/5 dysregulation in neuronal models within a described clinical mutation. GO-term enrichment for biological processes of ZARD NSCs reveals a significant downregulation of genes in pathways involved in BMP signaling regulation (FDR 1.0e-40), cellular response to BMP signaling (FDR 1.0e-22), and SMAD protein phosphorylation (FDR 1.0e-17) (**Figure 2A**). Notably *BMPR2*, the type II receptor essential for activating SMAD1/5 phosphorylation, is significantly downregulated in ZARD NSCs compared to controls with a (*p* value<0.0001) determined by RT-qPCR (**Figure 2B**). Interestingly, despite this downregulation, SMAD1 and SMAD 5 gene expression are slightly elevated (*p* value 0.0299 and 0.0320 respectively) determined by RT-qPCR of ZARD NSCs (**Figure 2C, D**). To directly assess SMAD1/5 activation, we performed p-SMAD1/5 immunostaining in ZARD NSCs. Quantitative image analysis revealed a significant decrease in phosphorylated SMAD1/5 in ZARD NSCs (*p* value 0.0002), confirming that SMAD1/5 phosphorylation is impaired providing a potential molecular mechanism for BMP-SMAD signaling cascade dysregulation (**Figure 2E, F)**. These findings demonstrate that the BMP-SMAD signaling cascade is dysregulated in ZARD NSCs, with reduced *BMPR2* expression leading to impaired SMAD1/5 phosphorylation and nuclear translocation. Here we propose that the *ZC4H2* transcription factor function is responsible for BMP-SMAD signaling cascade dysregulation with *BMPR2* being a DNA target of *ZC4H2*.

**Figure 2.**
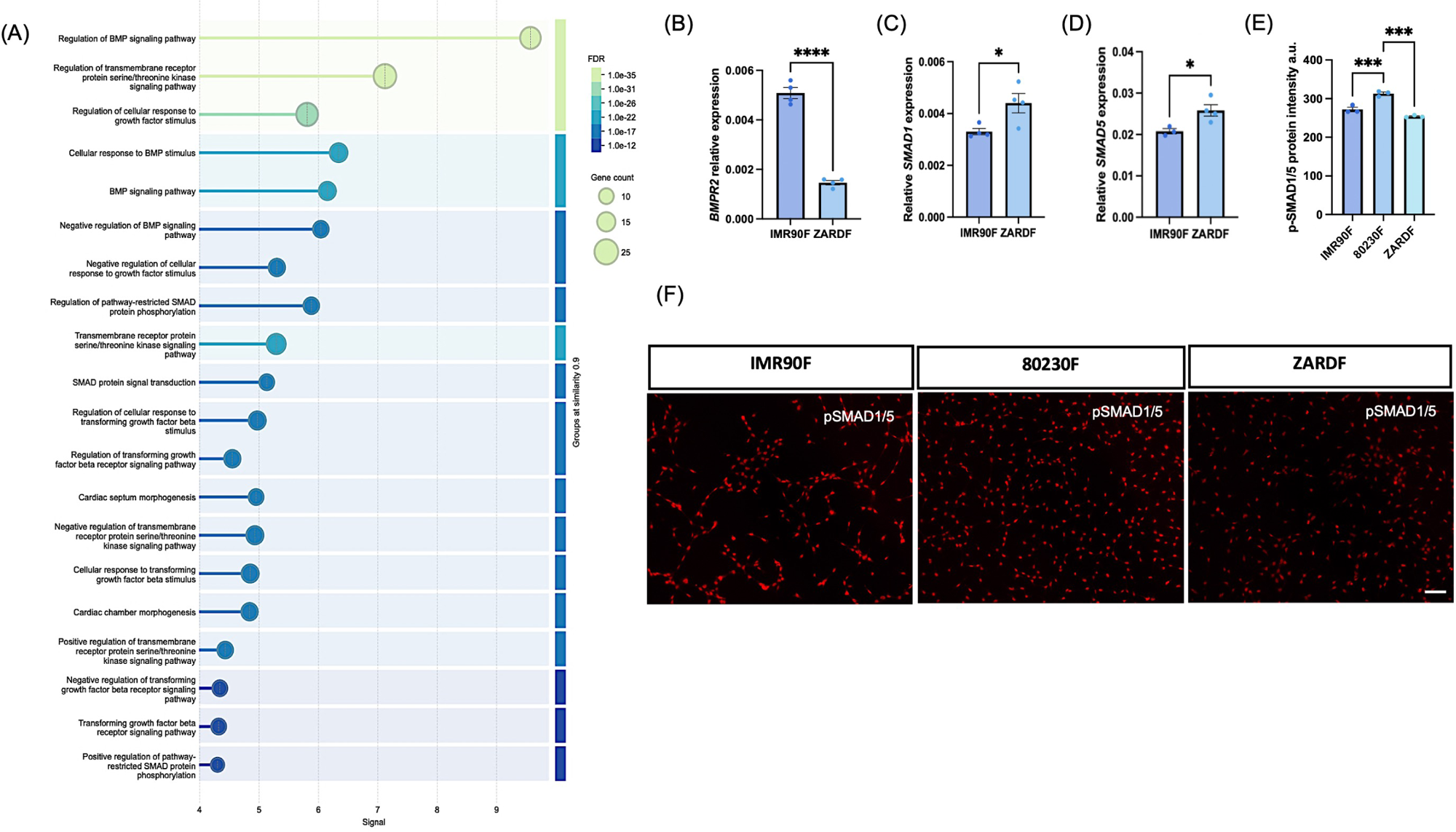
Dysregulation of BMP-SMAD signaling in ZARD NSCs. (A) Dot plot showing Gene Ontology (GO) Biological Process enrichment analysis of downregulated genes in ZARD NSCs (log2 fold change ≤ −2, adjusted *p* < 0.05). The x-axis represents enrichment significance (-log10 FDR). Dot size indicates the number of genes associated with each GO term, and dot color represents FDR. Enriched terms represent biological processes overrepresented among downregulated genes, including processes related to regulation of BMP signaling.(B-D) RT-qPCR analysis confirms reduced relative expression of *BMPR2* in ZARD NSCs compared to sex-matched control IMR90F NSCs *p* < 0.0001, followed by an increase in relative expression SMAD1 (p value 0.0312) and SMAD5 (*p* value 0.0320) in ZARDF NSCs (mean ± SEM, n = 3). Unpaired two-tailed t-test. (E) pSMAD1/5 (red) ICC of NSCs from control IMR90F, control 80230F, and ZARDF lines and subsequent quantification of protein intensity reveals significantly reduced pSMAD1/5 levels in ZARD NSCs compared to control lines. Mean ± SEM, n = 3 biological replicates. Statistical analysis was performed using one-way ANOVA with Dunnett’s multiple comparisons test, with 80230F as the reference group: 80230F vs. IMR90F, *p* value 0.0008; 80230F vs. ZARDF, *p* value 0.0001. Scale bar, 100 μm.

### shRNA-mediated knock down of *ZC4H2* recapitulates ZARD pathology

To validate the phenotypic effects of *ZC4H2* loss of function, control (80230F) iPSC derived NSCs were transfected with four short hairpin RNAs (shRNAs) targeting endogenous *ZC4H2* (**Figure 3A and B)**. The shRNA with the highest knockdown efficiency in NSCs was selected for further analysis. Two lead shRNAs (shRNA-01 and shRNA-02) significantly reduced *ZC4H2* mRNA expression by 50 and 52 % respectively determined by RT-qPCR (*p* value < 0.05) (**Figure 3B)**. Notably when looking at the differences in protein intensities post shRNA 01 and 02 treatment, we observed a ∼ 8% downregulation of *ZC4H2* protein with (*p* values of 0.0011 and 0.0021 respectively), showing a consistent reduction in *ZC4H2* protein levels post-knockdown. **(Figure 3C, 3D).** To assess whether shRNA-mediated knockdown replicates the dysregulation of genes critical for neurogenesis and the BMP-SMAD signaling cascade, additional RT-qPCR analysis was performed. Using shRNA-02 for subsequent experiments, results showed that a 52% reduction in *ZC4H2* resulted in a 50% decrease in *BMPR2* mRNA expression (*p* value 0.0164) and a 61% upregulation of *MAP2* mRNA (*p* value 0.0088) confirming these genes as *ZC4H2* targets (**Figure 3E, F**). Together, these findings demonstrate that *ZC4H2* knockdown disrupts key neurodevelopmental genes, including *MAP2* and *BMPR2*, and confirms the efficiency of shRNA-02 in targeting *ZC4H2* on and mRNA. These results demonstrate that *ZC4H2* LoF directly disrupts neurodevelopmental and BMP-SMAD signaling genes, recapitulating what was observed in variant NSCs.

**Figure 3:**
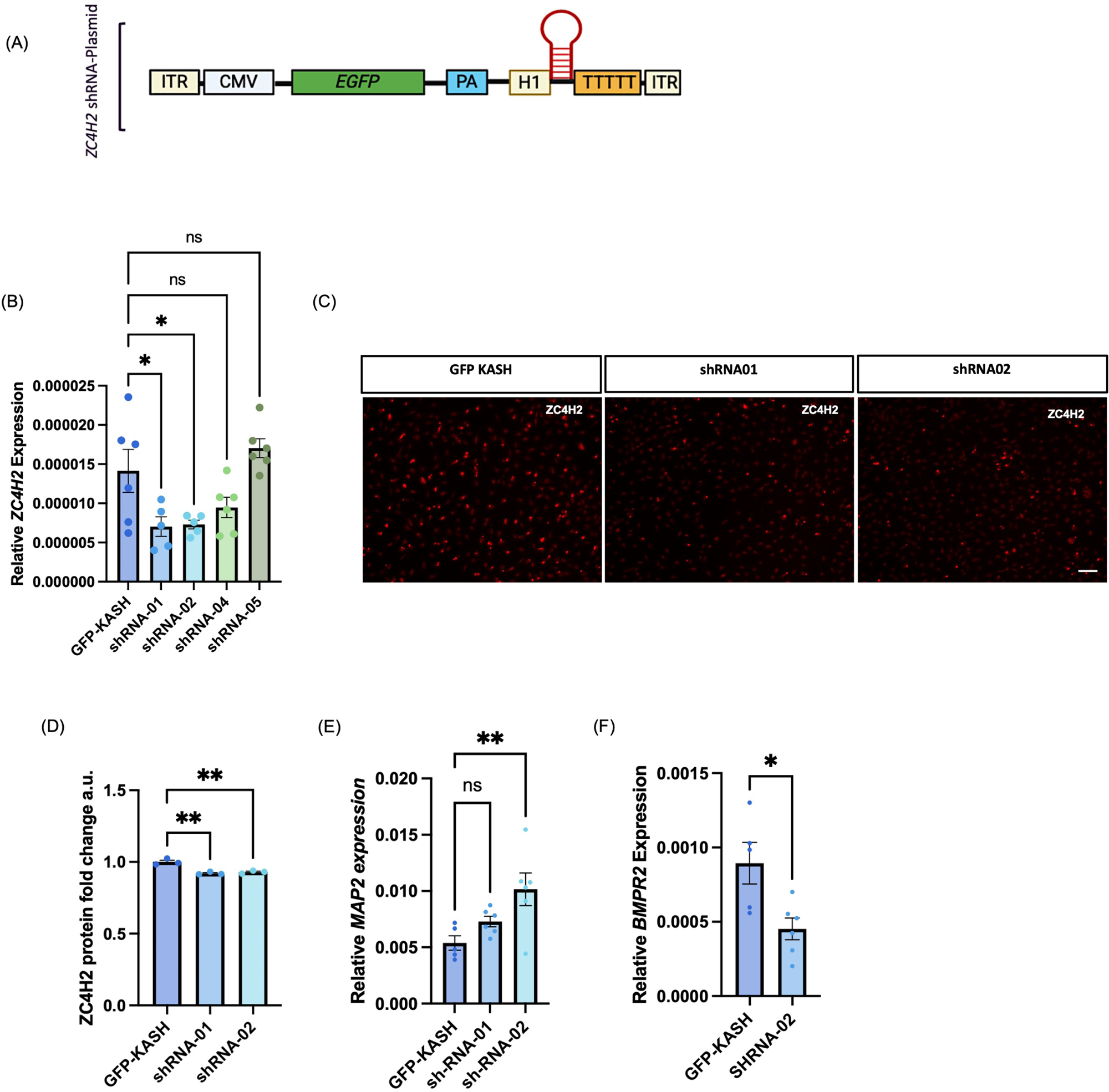
*ZC4H2* KD validates target genes. (A) Schematic of shRNA backbone that all shRNA were cloned into. (B) Relative *ZC4H2* expression in NSCs expressing GFP-KASH or *ZC4H2*-targeting shRNAs. Data is shown as mean ± SEM with individual replicates (n = 5-6). One-way ANOVA with Dunnett’s test was used to compare shRNA conditions to GFP-KASH control; *p* < 0.05, ns not significant. (C and D). *ZC4H2* (red) ICC of control 80230F NSCs transfected with GFP-KASH control or shRNA constructs (shRNA-01 and shRNA-02) followed by quantification of protein intensity reveals reduced ZC4H2 protein expression in shRNA-expressing NSCs compared to GFP-KASH controls. Mean ± SEM, *n* = 3 biological replicates. Statistical analysis was performed using one-way ANOVA with Dunnett’s multiple comparisons test: GFP-KASH vs. shRNA-01, *P* = 0.0011; GFP-KASH vs. shRNA-02, *p* value 0.0021. Scale bar, 100 μm. (E and F) Bar plot of relative *MAP2* and *BMPR2* gene expression in NSCs expressing GFP-KASH or ZC4H2-targeting shRNAs. Data are shown as mean ± SEM (n = 5-6). One-way ANOVA with Dunnett’s test; *p* < 0.01, ns not significant.

### Transcriptomic analysis of *ZC4H2* mutant CtOs

Following our initial characterization and mechanistic studies in NSCs, we extended our analysis to 3D cortical organoids (CtOs) derived from variant and control (80230F) female NSCs. RT-qPCR confirmed reduced *ZC4H2* expression in variant CtOs (*p* value 0.0133) (**Figure 4A**). Morphological analysis showed that ZARDF CtOs are significantly larger in size than 80230F CtOs over weeks 1–13 (extra sum-of-squares F test, F(2,276) = 303.9, *p* < 0.0001) (**Figure 4B, C**). Dissociated cell counts showed increased total cell number in variant CtOs as early as Day 18 in vitro (DIV) (*p* < 0.01), indicating that the abnormal NSCs proliferation observed in 2D cultures is maintained in 3D (**Figure 4D**). Consistent with our NSCs data, *BMPR2* and *SMAD5* expression were reduced in variant CtOs at 18 DIV (*p* < 0.05), confirming dysregulation of the BMP-SMAD signaling cascade within the 3D model (**Figure 4E, F**). To define how *ZC4H2* loss of function alters early neuronal transcriptional profiles, we performed bulk RNA-seq on female 80230F and ZARDF CtOs at 18 DIV and 46 DIV. Differential expression analysis revealed stage-specific transcriptional dysregulation, indicating that *ZC4H2* deficiency perturbs the temporal balance between progenitor maintenance, neuronal differentiation, and maturation. At 18 DIV, ZARD cortical organoids displayed widespread alterations in neuronal gene expression consistent with disrupted temporal coordination between neural stem cell maintenance and neuronal differentiation. Genes essential for neural stem cell maintenance and early neuronal structural development including *FAT3* (log2 - 8.00), *MAP1B* (log2 -8.11), *KIF1B* (log2 -6.69), *ARID1A* (log2 -6.52), and *RBBP4* (logFC -7.09) were significantly downregulated, whereas a subset of neuronal and synaptic genes typically expressed at later stages, such as *CBLN3* (log2 7.18), *SYTL4* (log2 5.83), and *CYFIP1* (log2 5.82) were upregulated **(Supplemental Table 3)**. These expression patterns highlight the presence of both reduced early neuronal maturation genes and increased late-stage neuronal markers within CtOs. By 46 DIV, this transcriptional dysregulation persisted. ZARD organoids exhibited coordinated downregulation of genes essential for the structural and functional maturation of neurons. Cytoskeletal and adhesion molecules including *TUBA1A* (log2 -9.65), *TUBB2B* (log2 -9.26), *MAP1B* (log2 -7.87), and *L1CAM* (log2 -8.27) key mediators of neurite outgrowth, axon guidance, and dendritic arborization, were markedly reduced, indicating compromised neuronal morphology and connectivity. Likewise, synaptic and signaling genes such as *STXBP1* (log2 -6.35), *NCAN* (log2 -8.36), *DPYSL2* (log2 -6.27), *DPYSL3* (log2 -8.20), *SCN9A* (log2 -6.03), and *MAPK10* (log2 -5.85) were downregulated, suggesting impaired synapse formation, neurotransmission, and electrophysiological maturation **(Supplemental Table 4)**. GO analysis of downregulated genes at 46 DIV revealed enrichment for neuron development, neuron projection development, and axon development, confirming that *ZC4H2* deficiency compromises neuronal maturation (**Figure 4G**). Collectively, these results propose a developmental model in which *ZC4H2* drives a premature shift toward neuronal fate but leads to persistent dysregulation of gene programs required for complete neuronal maturation, potentially disrupting the coordinated development of neural circuits underlying *ZC4H2*-related disorders, whose hallmark phenotypes include intellectual disability and seizures.

**Figure 4:**
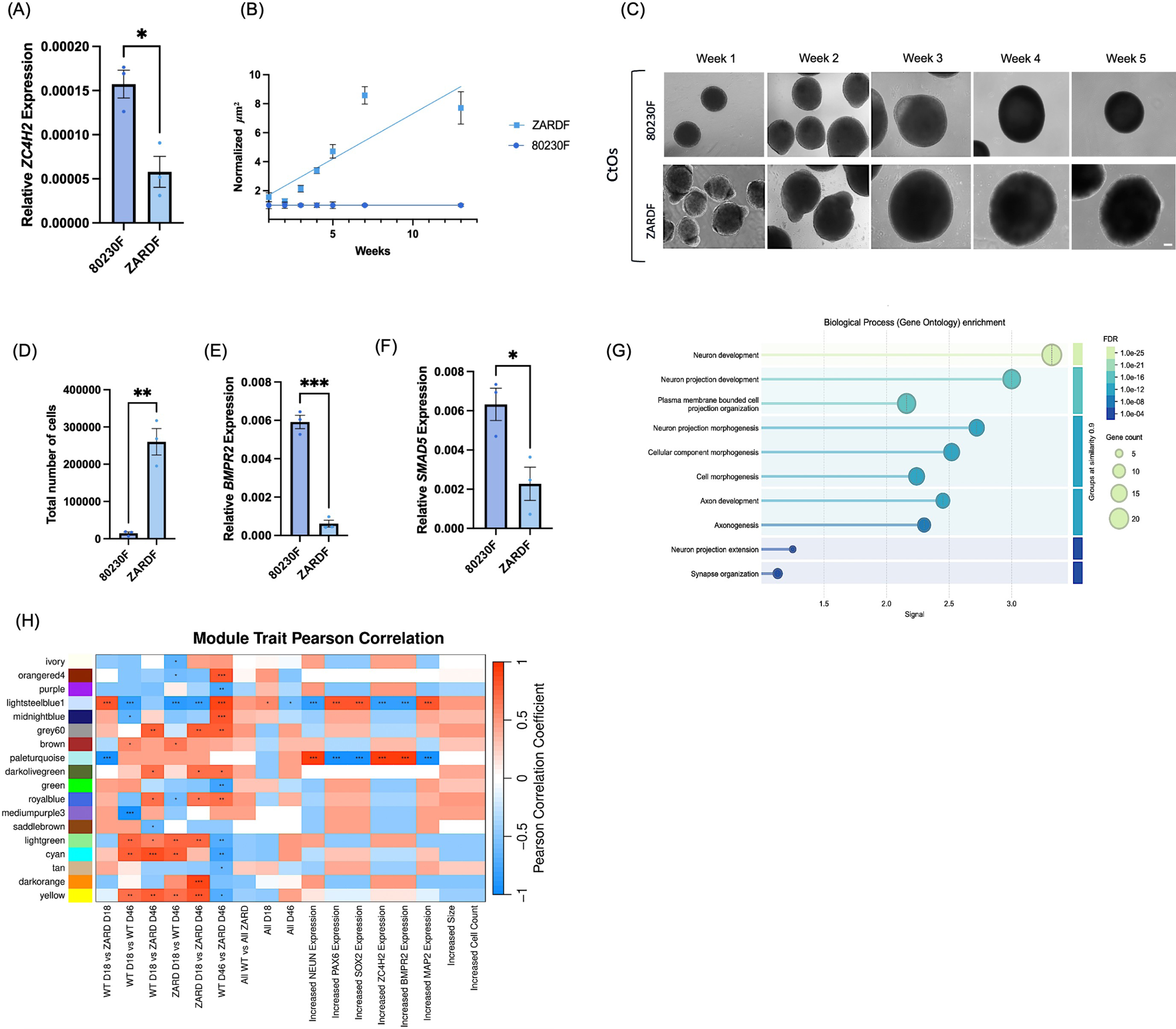
CtOs reveal temporal dysregulation of neurodevelopment. (A) RT-qPCR analysis showing significant downregulation of *ZC4H2* relative gene expression in variant CtOs compared to 80230F sex matched control. (B) Quantification of size changes in µm² (y-axis) were measured in ImageJ for a duration of 13 weeks (x-axis) N=20. All CtO area measurements are normalized to 80230F control. Data points represent mean ± SEM at each time point. Growth trajectories were analyzed using nonlinear regression with an exponential growth model, and curve comparisons were performed using an extra sum-of-squares F test, revealing significantly different growth dynamics between genotypes (F(2,276) = 303.9, *p* < 0.0001).(C) Representative image demonstrating CtO differences between control and variant CtOs over a span of 5 weeks. (D) CtO dissociation and quantification of single cells (y-axis) of control and variant CtOs (x-axis); N=3 per point. (E-F) RT-qPCR analysis showing significant downregulation of BMPR2 and SMAD5 relative gene expression (*p* value 0.0002 and 0.0265 respectively). (G) Dot plot showing GO Biological Process enrichment for downregulated pathways associated with the top ∼300 downregulated genes (log2 fold change ≥ 2, adjusted p <0.05) involved in neuron development. The x-axis represents enrichment significance (-log10 FDR). Dot size indicates the number of genes associated with each GO term, and dot color represents FDR. Enriched terms represent biological processes overrepresented among downregulated genes. (H) WGCNA module Pearson correlation heatmap. Red and blue shades denote the strength of positive and negative associations, respectively, between gene modules and specific experimental groups or traits. Positive correlations (red) link higher gene expression to the first group in a comparison, while negative correlations (blue) link it to the second group. All mRNA transfection experiments have an N between 3-6. Significance testing for quantification analyses was conducted by both unpaired t-test and one way ANOVA with statistical significance indicated as \*\*\**p*< 0.0001, \*\**p*< 0.001, \**p* < 0.05.

### Co-expression modules reveal temporal gene network dynamics distinguishing control and ZARD organoids

To further explore how these temporally misaligned transcriptional programs are organized at the network level, we performed Weighted Gene Co-expression Network Analysis (WGCNA) to identify transcriptional modules associated with genotype, developmental stage, and phenotypic traits. Six pairwise contrasts were examined to assess module trait relationships **(Figure 4H)**. When all wildtype (80230F) samples were compared to all ZARD mutants irrespective of stage, no module reached significance, indicating that genotype alone does not produce a global transcriptomic signature.

Because the developmental stage strongly influences global expression, genotype effects were interpreted only within stage-matched comparisons. Among the genotype-associated modules, the green and tan modules exhibited significant negative correlations with the 80230F D46 vs ZARD D46 contrast, indicating that genes within these networks are expressed at higher levels in 80230F organoids compared to ZARD. Functional annotation of the green module revealed enrichment for axon guidance pathways, supporting the interpretation that 80230F organoids possess a more active axonal wiring program at D46, whereas ZARD mutants may experience a deficit in this process, aligning with our GO data. The midnightblue module, enriched for axonogenesis and axonal-growth regulation, showed complementary trends of decreasing expression as 80230F CtOs matured but remained elevated relative to ZARD at 46 DIV indicating that axonogenesis networks fail to reach 80230F expression trajectories in ZARD organoids. Together, these findings define a transcriptional landscape shaped by developmental timing and genotype, in which WGCNA reveals that loss of *ZC4H2* function selectively perturbs gene networks associated with axonogenesis, offering mechanistic insight into the molecular basis of neuronal dysfunction in ZARD organoids.

### Reversal of ZARD related phenotypes via *ZC4H2* overexpression

To assess whether reintroducing *ZC4H2* can correct ZARD pathophysiology and rescue disease related phenotypes, we restored *ZC4H2* expression in human CtOs.A codon-optimized *ZC4H2* construct was packaged into AAV9 and transduced into DIV 100 cortical organoids (CtOs) at approximately 1 x 10^13^ viral genomes per 28 organoids **(Figure 5A, B)**. We generated CtOs from control 80230F and variant ZARD iPSC lines. Both lines were transduced with an AAV9 GFP-KASH control plasmid, whereas only ZARD CtOs were also transduced with an AAV9 *ZC4H2* overexpression plasmid using the same backbone. Four weeks post-transduction, CtOs transduced with *ZC4H2*-OE-vector shows a robust 2-fold increase in ZC4H2 protein expression (*p* value 0.0012) relative to the untreated ZARD CtOs **(Figure 5C,D; Supplemental Figure D).**To determine rescue of aberrant CtO size, treated CtOs size was measured as area (µm^2^) in ImageJ. CtO size dynamics differed by genotype and condition over time, with nonlinear regression revealing significant differences among WT (80230F-GFPKASH), and variant ZARDF-GFPKASH, and ZARDF-OE CtOs with ZARDF-OE showing a partial rescue of the ZARDF size phenotype as early as 3 weeks post-transduction **(Figure 5E)**. To further assess the cellular basis underlying changes in CtO size, dissociated single cells were quantified across control and variant CtOs. ZARD-GFPKASH CtOs yielded a significantly higher number of dissociated cells compared to control CtOs, whereas ZC4H2 overexpression resulted in a significant reduction in single-cell counts, restoring values to levels comparable to control (one-way ANOVA with multiple comparisons). These findings are consistent with the partial rescue of CtO size observed following ZC4H2 overexpression. **(Figure 5F).** Visual inspection of CtOs four weeks post-transduction revealed no obvious morphological features indicative of toxicity or increased cell death **(Figure 5G).** RT-qPCR analysis demonstrated rescue of *ZC4H2* gene by 350,000 fold when compared to untreated ZARD CtOs *p* value< 0.0001. When looking at changes in target *ZC4H2* gene *BMPR2* expression RT-qPCR analysis determined a 4-fold increase in *BMPR2* gene expression post *ZC4H2* OE treatment (*p* value 0.0372) **(Figure 5H, I)**. Given that *BMPR2* gene expression is rescued by 4-fold post AAV treatment, we wanted to determine whether this leads to rescue of SMAD1/5 phosphorylation. Immunostaining and quantification show a ∼4-fold increase in SMAD1/5 phosphorylation in AAV treated CtOs when compared to ZARD-GFP-KASH controls p-value (0.0486). When comparing UT-ZARD vs OE-ZARD CtOs we see a clear trend towards an increase in pSMAD1/5 phosphorylation, yet significance was not reached p-value (0.1346) **(Figure 5J,K, Supplemental Figure E-F)**.Previous reports have demonstrated *ZC4H2* protein to be important for synaptogenesis within excitatory neuron post synaptic density (PSD), with the loss of *ZC4H2* protein correlated with increased dendritic spine density compared to controls. MAP2 protein expression is essential for neurite outgrowth, dendritic branching, and neuronal maturation; therefore, maintaining appropriate MAP2 levels during early neurodevelopment may help prevent aberrant neural development and support normal neuron formation. In ZARD cortical organoids (CtOs), *MAP2* gene expression increased significantly by 18 DIV post-aggregation (**Supplemental Figure G**). Notably, *ZC4H2* OE (*ZC4H2*-OE) reduced MAP2 protein levels in treated ZARD CtOs compared with UT-ZARD CtOs by by approximately 48% relative to control (mean fold change = 0.52; *p* value 0.0183 (**Supplemental Figure H,I).**

**Figure 5:**
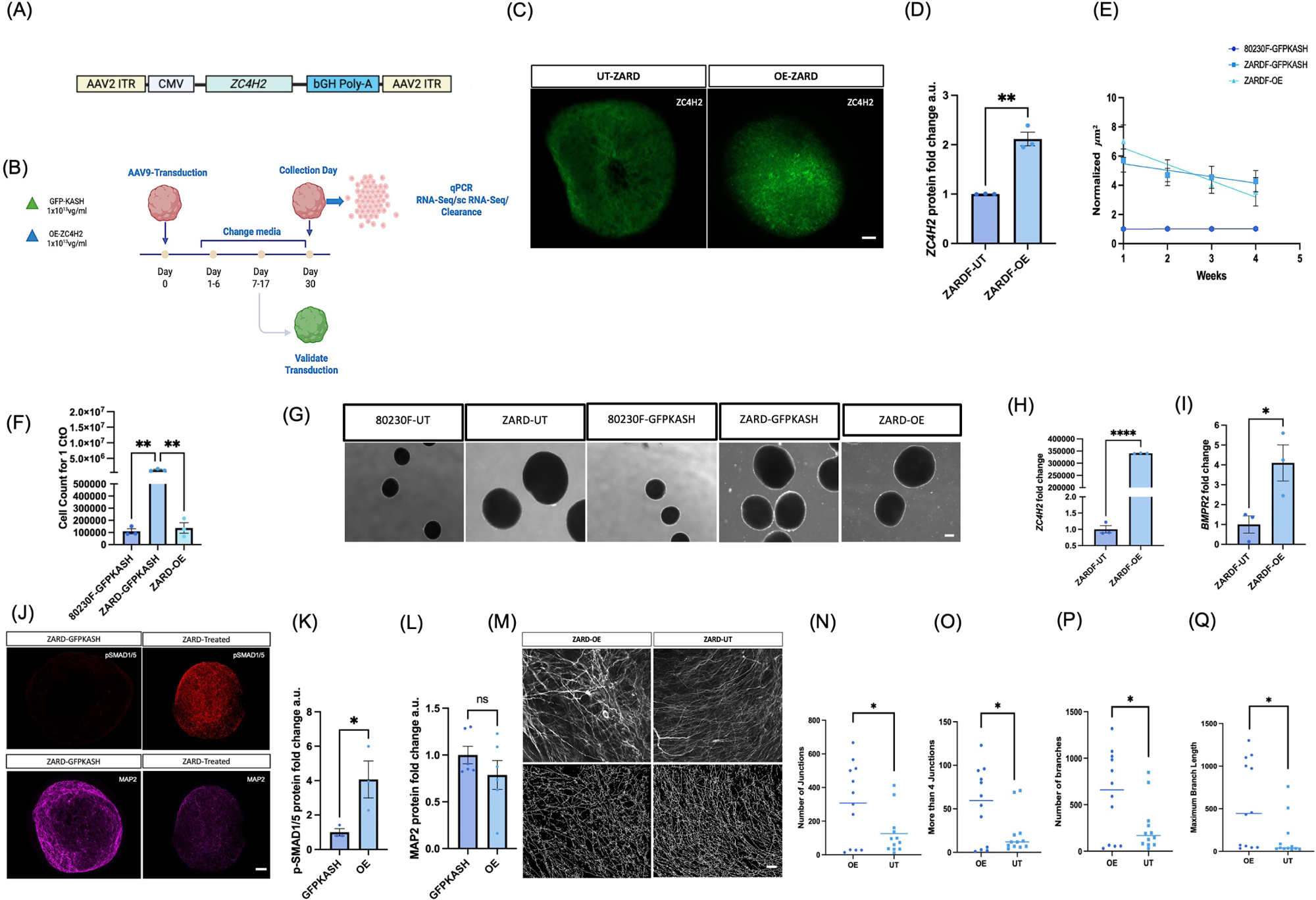
Rescue of ZARD Cellular and Molecular Phenotypes via AAV9-Mediated Overexpression in CtOs. (A) Schematic of the AAV9 OE construct used for gene delivery. (B) Schematic diagram of AAV9 transduction protocol applied to DIV 100 CtOs at a dose of 1 x 10¹³ viral genomes per 28 organoids. (C) Representative images of ICC-stained control untreated variant and OE treated variant CtOs for ZC4H2 protein expression post 4-week transduction. Image taken at 10X with Scale bar, 100 μm. (D) Cleared and immunostained CtOs expressing the OE construct show increased ZC4H2 protein levels compared to control, as quantified by macro-based fluorescence intensity analysis. Data are presented as mean ± SEM from three biological replicates. significance was assessed via an unpaired two-tailed Student’s t-test(*p* value 0.0012).(E) Quantification of size changes in µm² (y-axis) for a duration of 4 weeks post transduction of AAV9 (x-axis) N=15. CtO growth dynamics differed significantly by genotype and condition, as determined by nonlinear regression with extra sum-of-squares F tests, revealing differences among 80230F-GFPKASH, ZARDF-GFPKASH, and ZARDF-OE CtOs (F(4,174) = 313.0, *p* < 0.0001). ZARDF-GFPKASH CtOs differed from 80230F-GFPKASH controls (F(2,116) = 838.9, P < 0.0001) and ZARDF-OE CtOs (F(2,116) = 11.56, *p* < 0.0001), with ZARDF-OE CtOs showing partial rescue of the ZARDF size phenotype. (F) CtO dissociation and quantification of single cells (y-axis) from control and variant CtOs (x-axis); n = 3 biological replicates per condition. ZARD-GFPKASH CtOs yield a significantly higher number of dissociated single cells compared to control 80230F-GFPKASH CtOs, whereas ZARD-OE CtOs show reduced single-cell counts relative to ZARD-GFPKASH and are not significantly different from control. Data are presented as mean ± SEM. Statistical analysis was performed using one-way ANOVA with Dunnett’s multiple comparisons test: ZARD-GFPKASH vs. 80230F-GFPKASH, p value 0.0010; ZARD-OE vs. ZARD-GFPKASH, *p* value 0.0012; ZARD-OE vs. 80230F-GFPKASH, not significant (*p* value 0.9826).(G) Representative brightfield images showing changes in CtO size at week 4 post-transduction. (H) RT-qPCR analysis revealed a ∼350,000-fold change in *ZC4H2* gene expression in ZARDF CtOs treated with OE plasmid (*p* value 0.0001). (I) RT-qPCR revealed a 4-fold increase in BMPR2 expression (*p* value 0.0372) unpaired t-test. (J) Representative Z-stack images of immuno-stained treated CtOs with pan-neural marker MAP2 and phosphorylated SMAD1/5 protein expression at 10X with scale bar, 100 μm. (K and L) Quantification of MAP2 and PSMAD1/5 protein intensity were quantified using a FIJI macro-N=3-6 organoids. unpaired t-test revealed a significant upregulation of SMAD1/5 phosphorylation post AAV9 treatment. MAP2 protein level were reduced but trend did not reach significance. (M) Representative 40X images of treated vs untreated cortical organoids alongside Sholl and skeleton generated images mapping out neuronal projections; scale bar, 100 μm. (N-Q) Sholl/skeleton analysis revealed an increase in junction number, branch number, and branch length post AAV9 treatment. Analysis was done on 3 biological replicates with 4 images taken at 40X per biological replicate. Significance was determined using an unpaired t-test \*\*\**p*< 0.0001, \*\**p*< 0.001 \**p* < 0.05. All photos were taken using the zeiss brightfield microscope and deconvoluted for further analysis.

When compared with the GFP-KASH control, MAP2 protein expression showed a clear downward trend, though this difference did not reach statistical significance **(Figure 5L; Supplemental Figure J).** *ZC4H2* plays a critical role in regulating neuronal complexity, including dendritic spine density, dendritic branching, branch length, and synapse number, thereby maintaining the structural and functional integrity of neurons. To examine the effects of *ZC4H2* restoration on neuronal complexity we mapped out the neuronal projections of GFP and *ZC4H2-OE* transduction CtOs via Sholl analysis. Sholl analysis revealed that ZARD neurons exhibit reduced complexity, characterized by a decrease in the number of dendritic branches, branch length, and junctions. Interestingly, 4 weeks post transduction, aberrant neuron morphology was rescued. Specifically, quantitative analysis paired with Sholl analysis revealed that *ZC4H2* overexpression increases the number of neuronal branches, branch length, and junction points overall rescuing neuronal complexity **(Figure 5M-Q)**. This study is limited to morphology studies; additional analysis needs to be done to access functional rescue of synaptic activity. Overall, these findings support codon-optimized *ZC4H2* delivery via AAV9 can rescue protein expression of MAP2, PSMAD1/5, and neuronal complexity highlighting *ZC4H2*-OE as a therapeutic strategy for ZARD.

## Discussion

In this study, we established and characterized female patient-derived NSCs and CtOs carrying the variant ARR[HG19] XQ11.2(64,170,839–64,414,573) X1 to investigate ZARD pathology. In ZARD NSCs, transcriptomic analysis indicated widespread dysregulation of gene programs involved in stem cell maintenance and neuronal differentiation. Unlike prior *Zc4h2* knockout mouse reports of inhibited NSCs proliferation (8), we observed accelerated NSCs proliferation, alongside premature neuronal fate. Previous reports for iPSC-derived CtOs have characterized 18 DIV to correspond to the fetal cortex 6-8 weeks post conception representing initial neural stem cell proliferation, and early neural induction while day 46 DIV, organoids represent early neuronal differentiation and development mapping to approximately 10-12 weeks post conception(12,13). Because *ZC4H2* is proposed to be an important gene for regulating early neurodevelopment, these time points allowed us to understand the dysregulation of early neurodevelopment in a *ZC4H2*-deficient environment. Transcriptomic analysis of these two developmental time points suggests temporally dysregulated NSC and neuronal programs promoting premature neuron differentiation without proper structural neuronal maturation.

Similar abnormalities in developmental timing have been described in Fragile X syndrome, where increased progenitor proliferation occurs alongside impaired neuronal maturation(14). In ZARD, elevated early neurogenic signatures together with reduced expression of neuronal structural maturation genes such as *MAP1B*, *KIF1B*, and *FAT3* within ZARD NSCs and CtOs suggest a disruption in the coordination between early neurogenesis and later stages of neuronal development. Such timing defects may hinder neuronal integration into neuronal networks and represent a mechanism shared across multiple neurodevelopmental disorders. Additional studies need to be done to understand how this dysregulation affects neuronal network excitability. Morphological analysis of neuron complexity in AAV9 *ZC4H2* transduced CtOs demonstrated improved neuronal complexity when compared to untreated ZARD CtOs. Further studies need to be done to validate these findings alongside WT neurons. Size analysis showed ZARD organoids were larger than controls reflecting the heightened NSCs proliferation seen in variant NSCs, although macrocephaly has yet to be clinically described within this variant. Mechanistically, our data implicates the BMP-SMAD signaling cascade as a central pathway disrupted in variant NSCs and CtOs. Prior work in Xenopus showed that *ZC4H2* stabilizes SMAD1/5 protein by preventing its ubiquitination and subsequent degradation, providing stability for the BMP-SMAD signaling. Consistent with this model, we found that the 244 kb ZARD mutation is associated with depletion of BMPR2 and reduced p-SMAD1/5 in NSCs and CtOs, because BMPR2 phosphorylates BMPR1, which then phosphorylates SMAD1/5, reduced BMPR2 likely limits SMAD1/5 activation; unphosphorylated SMAD1/5 cannot form heteromeric complexes with SMAD4, blocking nuclear translocation and target-gene activation disrupting the activation of the BMP-SMAD signaling cascade. In mice, inhibition of BMP-SMAD signaling via noggin overexpression or BMPR2 ablation promotes NSCs cell cycle reentry and accelerates neurogenesis without depleting the stem cell pool, which aligns with our phenotype(15).

Here we show that *ZC4H2* knockdown decreased BMPR2 mRNA levels supporting that *ZC4H2* regulates *BMPR2* at the transcript level and establishes BMP-SMAD dysregulation as a mechanism underlying this ZARD variant. We demonstrated that delivery of AAV9 ZC4H2-OE at 1×10^13^vg/ml greatly elevated *ZC4H2* mRNA expression, yet this only resulted in a 2-fold increase in protein expression indicating a ceiling in the amount of ZC4H2 protein being expressed. Here we describe that 2-fold increase in ZC4H2 protein expression is sufficient to rescue *BMPR2* gene expression, p-SMAD1/5 phosphorylation, normalizes CtOs size, and increases neuron complexity when compared to untreated ZARD controls demonstrating therapeutic potential for *ZC4H2* reinstatement applicable to both male and female ZARD patients.

Overall, our results motivate two hypotheses about pathophysiology. First, loss of *ZC4H2* accelerates neuronal fate while hindering neuron development and maturation. Prior studies have identified *ZC4H2* as a postsynaptic regulator that interacts with AMPARs and controls their ubiquitination to maintain receptor stability; in *ZC4H2* conditional knockouts, AMPARs accumulate at excitatory synapses, synaptic transmission is heightened, and plasticity is disrupted(16). Additionally, *ZC4H2*-deficient excitatory neurons exhibit increased dendritic spine density, enhanced dendritic branching, and elongated apical dendrites, highlighting its role in regulating excitatory neuron arborization. Notably, when comparing neuronal morphology between *ZC4H2* overexpression and untreated controls, we observed an increase in neuronal complexity, highlighting the importance of *ZC4H2* in regulating neuronal arborization. In our variant CtOs, we observed dysregulation of genes consistent with premature neuronal yet temporally dysregulated maturation when compared to WT.

A second hypothesis is that the observed maturation deficits may stem from reduced BMP-SMAD signaling observed in LoF model. Previous reports have demonstrated that SMAD1 regulates excitatory neuron intervention on GABAergic PV-interneuron, through the regulation of gene programs required for excitatory synapse formation and perineural net formation(17). It was demonstrated that loss of SMAD 1 resulted in hyperactivity due to the reduced functional recruitment of inhibitory neurons. They also found that SMAD1 loss affects excitatory-inhibition neuronal balance. Hyperactivity has been reported in mouse models yet whether the loss of *ZC4H2* is associated with excitatory-inhibitory neuronal imbalances remains elusive. This study is limited to female patient-derived model of a single variant, emphasizing the need to characterize and test therapeutic strategies across additional ZARD genotypes and developmental windows to define shared versus variant-specific mechanisms and responses to *ZC4H2* restoration. Even so, our findings identify BMP-SMAD dysregulation as a feature of ZARD in patient-derived models and provide proof-of-concept that restoring *ZC4H2* by overexpression of *ZC4H2* can rescue molecular, cellular, and growth phenotypes.

## Methods

### Human Subjects

Human iPSC lines derived from a female patient diagnosed with ZC4H2-associated rare disorder (Wieacker-Wolff syndrome, female-restricted) were obtained from the Coriell Institute for Medical Research Biobank (Catalog ID: GM28603).

### hiPSCs derived NSCs Generation and Culture

iPSCs were enzymatically dissociated using 0.75x TrypLE Select in PBS (phosphate buffered saline) and seeded at a density of 3×10⁶ cells/ per well in AggreWell 800 plates with embryoid body (EB) Medium (stem cell technologies) supplemented with Y27632 (10 μM). To aggregate iPSCs into EB, iPSCs were centrifugated at (1300xg, 5 min, room temperature (RT). EB were cultured overnight at 37 °C The following day, EBs were transferred to low-attachment plates in EB Medium containing Noggin (500 ng/mL) and SB431542 (10 μM), with media changes every other day. On Day 5, EBs were plated onto Growth Factor Reduced Matrigel-coated plates (1:20 dilution in cold DMEM/F12) and maintained in induction supplemented with Noggin and SB431542. Beginning Day 6, culture medium was switched to Neural Progenitor Medium, refreshed every other day. Around Day 11-12, neural rosettes were dissociated with accutase (3-4 min, 37 °C) and seeded at 50,000-100,000 cells/cm² on Poly-L-Ornithine/Laminin-coated flasks in NSC Medium Medium) supplemented with Y27632 (10 μM). NSCs were passaged at 80% confluency and cryopreserved in NSC Freeze Medium.

### NSC to CtO derivation

NSCs were enzymatically dissociated with acutase and counted prior to aggregation. (3×10⁶ cells per well) were resuspended in AggreWell EB Medium supplemented with 10 μM Y27632 and plated into pre-treated AggreWell 800 plates primed with DMEM/F12 and anti-adherence rinsing solution. Plates were centrifuged at 100xg for 3 minutes to ensure even cell distribution, then incubated overnight at 37 °C for spheroid formation. On Day 1, EB were gently dislodged from AggreWell wells, filtered through a 37 μm reversible strainer, and transferred to ultra-low attachment 6-well plates, splitting each AggreWell well into three wells containing 3 mL of NSC Medium each. Culture media were changed according to the following schedule: NSC Medium daily from Days 2-6; every other day from Days 7-17. LC Medium every other day from Days 18-45; BrainPhys+ Medium (stem cell technologies) every other day from Days 46-90; and BrainPhys+ Medium three times weekly after Day 90. During media changes, plates were tilted carefully to collect organoids without disturbance, maintaining approximately 20 organoids per well from Day 18 onward.

### Real-time quantitative PCR

Total RNA was extracted using Direct-zol RNA miniprep according to the manufacturer’s recommendations ZYMO Research kit. 250ng of RNA were reverse transcribed into cDNA using the iScript™ cDNA Synthesis Kit, 500 x 20 µl rxns #1708891BUN. Real time RT-qPCR was performed using Applied Biosystems™ PowerUp™ SYBR™ Green Master Mix for RT-qPCR Cat: A25742, and the QuantStudio 6 Flex System. The thermal cycling protocol consisted of an initial hold stage at 50 °C for 2 min with a ramp rate of 1.9 °C/sec, followed by a second hold stage at 95 °C for 2 min with the same ramp rate. Then, during the PCR stage, samples were heated to 95 °C for 1 sec at a ramp rate of 1.9 °C/sec, then incubated at 60 °C for 30 sec at a ramp rate of 1.6 °C/s. The melt curve stage comprised followed with 95 °C for 15 s at 1.9 °C/s, 60 °C for 1 min at 1.6 °C/s, and 95 °C for 15 s at a ramp rate of 0.05 °C/s.All RT-qPCR analyses were conducted using RNA extracted from at least 3 independent biological replicates per subject/condition, with 3 technical replicates for each probe set/sample combination. Normalization was achieved using GAPDH as an endogenous control gene, and relative expression was calculated using the comparative ΔΔCt method. The following RT-qPCR primers were used are in (**Supplemental table 1**).

### CtO size and total cell count analysis

To assess organoid size, 10-20 images were captured per organoid at 4x magnification. Size was quantified by applying an intensity threshold of 0-55 in ImageJ and calculating the area in (µm²) using ImageJ. For each experiment and time point, size measurements were normalized to the mean value of the wildtype control 80230F. Time-course data were analyzed by nonlinear regression using an exponential growth model with least-squares fitting. Differences in size dynamics among groups were assessed using extra sum-of-squares F tests. Statistical significance was defined as *p* < 0.05. For cell count analysis. Organoids (3 per tube, n = 3 tubes) were washed twice in PBS and sonicated for 5 s in 0.1 mL PBS using a probe sonicator. Cell counts were obtained using an automated Cell Countess.

### Fixation and Immunocytochemical staining of NSCs

NSC were seeded 1 day prior to fixation at a density of 50,000 cells/cm². Media were removed and wells rinsed gently with PBS. Cells were fixed with 1 mL of 4% paraformaldehyde (PFA) per well at RT for 15 minutes. Fixative was removed, and wells were washed with 1x Immunocytochemistry (ICC) Wash Buffer (prepared by diluting 10x stock 1:10 with molecular-grade water and stored at 4 °C). Cells were permeabilized with 1 mL of 0.1% Triton X-100 in PBS per well for 15 minutes at RT. Blocking was performed using 1 mL of 3% Bovine Serum Albumin (BSA) in PBS per well for 1 hour at RT. Primary antibodies were diluted in the blocking solution and applied at 500µl per well; control wells received blocking buffer only. Plates were incubated overnight at 4 °C. Following primary antibody incubation, wells were washed three times with ICC Wash Buffer, 5 minutes each at RT. Secondary antibodies were diluted in blocking solution and applied at 500 μL per well, with plates wrapped in foil and incubated for 1 hour at room temperature on a shaker (rpm) in the dark. Wells were washed three more times with ICC Wash Buffer, then stained with 500 μL of Hoechst 33342 (1:2000 dilution in PBS) per well for 5 minutes in the dark at RT with no agitation. Cells were washed once more and stored in 1 mL ICC Wash Buffer at 4 °C, protected from light, for up to one week. Antibodies and concentrations: MAP2 1:750; SOX2 1:400; TUJ1 1:500; NESTIN: 1:100 pSMAD1/5 1:200; GFAP: 1:200

### Fixation, Immunocytochemistry, and Optical Clearing of CtOs

Organoids were fixed overnight at 4 °C in 4% (PFA) and 8% sucrose in phosphate-buffered saline (PBS). After fixation, samples were washed three times for 1 hour each at room temperature (RT) with gentle rocking in PBS, then transferred to 48-well plates. Using the CytoVista Tissue Clearing Kit (Thermo Fisher Scientific) organoids were incubated for 30 minutes at RT in Cytovista™ Antibody Penetration Buffer containing 0.2% Triton X-100, 0.3 M glycine, and 20% DMSO in PBS to improve antibody penetration. Samples were washed twice for 10 minutes each at RT in PBS, then blocked with Cytovista™ Blocking Buffer for 2 hours at 37 °C with gentle agitation (100 rpm). Primary antibodies diluted in Cytovista™ Antibody Dilution Buffer were applied overnight at 37 °C with gentle shaking (100 rpm). After incubation, samples were washed five times for 5 minutes each at RT in Cytovista™ Washing Buffer (1x), which contains 0.2% Tween-20, 10 µg/mL heparin, and 1% BSA. Secondary antibodies and nuclear dyes were applied overnight (∼16 hours) at 37 °C with gentle agitation (rpm). Hoechst 33342 was used at 1:1000 dilution in PBS for 1 hour at 37 °C, followed by two washes of 10 minutes each at RT in PBS. For optical clearing, organoids were transferred to glass-bottom well plates and incubated overnight (∼16 hours) at RT in 1 mL CytoVista™ Tissue Clearing Reagent with gentle shaking (100 rpm). Cleared samples were stored at 4 °C in the dark until imaging. For mounting, cleared organoids were placed on Superfrost Plus charged glass slides with four strips of (scotch permanent double-sided tape 0.5 in. x 500 in., with a 1-in. core) as spacers to prevent compression. A PAP pen and dried prior to adding approximately 75-80 µL of clearing reagent with the organoid. Samples were cover slipped with #1.5 coverslips, sealed, and stored at 4 °C in the dark until imaging. Fluorescent imaging was performed using an Evos bright field microscope. All images were taken using the apotome at 10X magnification. Post- image processing included Apotome deconvolution (FIJI version: 2.16.0/1.54p) for Z-Stack projections (18,19). Antibodies and concentrations: MAP2 1:750; ZC4H2 1:100; pSMAD1/5: 1:200

### Proliferation Assays

Cells were pulse-labeled with 10 µM EdU for 2.5 h (2 µL of 10 mM stock per 2 mL medium) using the Click-iT™ EdU Flow Cytometry Assay Kit (Thermo Fisher Scientific), then dissociated with Accutase, washed in 1% BSA/PBS, and pelleted at 500 x g for 5 min. Pellets were fixed for 15 min at 25 °C in the kit fixative, rinsed with 1% BSA/PBS, and permeabilized for 15 min in 1x Click-iT saponin-based permeabilization/wash reagent. The Click-iT reaction cocktail (prepared per manufacturer’s instructions with freshly made 1x buffer additive; 10x stock diluted 1:10 in H₂O) was added and samples incubated 30 min at RT protected from light with gentle mixing, followed by a wash in 1x permeabilization/wash reagent and resuspension in PBS + 10% FBS. For DNA content, cells were incubated in PBS/0.1% Triton X-100/100 µg/mL RNase A containing 20 µg/mL propidium iodide, then kept on ice and protected from light for acquisition. Per sample, 2 x 10⁵-1 x 10⁶ cells were analyzed by flow cytometry; EdU-positive cells were quantified in the channel appropriate for the kit fluorophore.

### Cloning of sgRNAs

For transient transfection experiments and transduction experiments, sgRNAs were cloned into a sgRNA expression vector (Addgene plasmid # 180280) following previously published protocol. In parallel, the CRISPR activation effector plasmid was prepared (Addgene plasmid #176269). All constructs were sequence-confirmed by Sanger sequencing (Genewiz, Inc, South Plainfield, NJ, USA) and chromatograms were analyzed using SnapGene software (from GSL Biotech; available at snapgene.com).

### *ZC4H2* Over expression

Codon optimized *ZC4H2* over expression construct was made using IDT. EcoRI and AgeI restriction enzymes allowed for the insertion of *ZC4H2* into a pAAV-EGFP-KASH backbone.

### *ZC4H2* Knockdown shRNA generation

ShRNA knockdown oligos were generated using the GPP Web Portal https://portals.broadinstitute.org/gpp. NheI and BamHI were used to clone shRNA oligos into the pAAV-EGFP-KASH backbone.

### Neuronal Network Analysis

Following optical clearing of cerebral organoids, MAP2-stained images were acquired at 20x magnification from three organoids per condition (OE and UT), with four randomly selected fields per organoid captured, excluding regions corresponding to organoid edges. To assess neuronal network complexity, segmentation and quantification of neuronal structures were performed in Fiji/ImageJ (version 1.54, NIH, Bethesda, MD, USA) using a custom macro. Briefly, local thresholding was applied to raw images using the Phansalkar method (radius = 14 pixels), and small objects (<15 pixels) were removed to exclude debris. The processed images were subsequently analyzed with the Analyze Skeleton (2D/3D) plugin to quantify neurite morphology(18,20–22). Extracted parameters included the number of branches, shortest path length, number of junctions, and total neurite length. For each image, the main trunk (largest connected neurite structure) was selected to represent the central network of the organoid(23). Comparative analysis was then performed on the extracted main trees across conditions. Parameters included the total number of junctions, the frequency of higher-order junctions (≥4 branches per junction), the total number of branches and the maximum neurite length within the main tree.

### RNA-seq Bioinformatics Analysis

The RNA-seq DEG and WGCNA were conducted as previously described(4). Briefly, raw sequence data was trimmed using Trim Galore (v0.6.10), aligned to the hg38 genome using STAR3 (v2.7.11a) with the hg38 NCBI RefSeq GTF file for annotation, indexed using Samtools5 (v1.21), and subject to quality control using MultiQC6 (v1.29)(24–27).Gene names from raw count data were mapped to gene names from the HUGO Gene Nomenclature Committee, after which the counts were read into a Jupyter Notebook7 (v7.4.0) running R8 (v4.4.2) via IRkernel9 (v1.3.2). The data were then normalized and filtered using the standard method described by Chen et al. 2016 (28), and subsequently by genes with CPM > 1 in at least 15% of samples, resulting in a final set comprising any gene that satisfied either criterion. To evaluate the effects of genotype (ZARD vs. 80230F) and time point (D18 vs. D46), all pairwise combinations of sample conditions were subject to DEG analysis using limma11 (3.62.2) in conjunction with edgeR12 (v4.4.2), yielding the following comparisons: 80230F D18 vs. ZARD D18 80230F D18 vs. 80230F D46, 80230F D18 vs. ZARD D46, ZARD D18 vs. 80230F D46, ZARD D18 vs. ZARD D46, and 80230F D46 vs. ZARD D46 (n = 3 per group) (29–33). Heatmaps were generated from the normalized counts of all statistically significant DEGs (adjusted p-value < 0.05). Euclidean distance was used for hierarchical clustering. We also performed co-expression analysis with WGCNA13 (v1.73). After soft thresholding, we built block wise modules (power = 25) and computed module eigengenes to capture each module’s overall expression pattern(34). We used Pearson correlation to correlate modules to sample traits. GO was conducted for DEG and WGCNA results. For the DEG analysis, we filtered genes to an adjusted p-value < 0.05 before submitting them, whereas for WGCNA we submitted all genes belonging to each module to the following Enrichr15 (v3.4) databases: GO Biological Process 2025, GO Cellular Component 2025, GO Molecular Function 2025, KEGG 2021 Human, Panther 2016, Reactome 2022, and RNA-seq Disease Gene and Drug Signatures from GEO.Plots associated with the DEG and WGCNA results were created using the following packages: ggplot216 (v3.5.2), Heatmap software(35–37).

## Acknowledgement

This work was supported by a research grant from the University of Pennsylvania Orphan Disease Center in partnership with the ZC4H2 Research Foundation (MDBR-23-036-ZC4H2 to JH). YP was supported by the California Institute for Regenerative Medicine (CIRM) Ph.D. Fellowship (Award EDUC4-12792) through the UC Davis Cell and Gene Therapy Center. The contents of this publication are solely the responsibility of the authors and do not necessarily represent the official views of CIRM or any other agency of the State of California. The authors declare that they have no conflict of interest.

## Figure legends

**Supplemental Figure 1:** (A) Growth curve of single-cell NSC counts (y-axis) over a 4-day period (x-axis). Nonlinear regression analysis revealed that IMR90F and ZARDF exhibited significantly different growth dynamics over time (extra sum-of-squares F test, F(2,20) = 5.20, *p* value 0.0152). (B) Volcano plot showing differential gene expression between conditions. Log₂ fold change is plotted against −log₁₀ p-value. Significantly upregulated (red) and downregulated (blue) genes are highlighted, with the top five most upregulated and downregulated genes labeled.(C) Quantification of ICC MAP2–stained NSCs. MAP2 protein expression did not reach statistical significance (unpaired two-tailed t-test, *p* value 0.1344).(D) Representative snapshots of treated AAV9-OE and untreated (UT) ZARDF CtOs at 4 weeks post-transduction (10x). Scale bar, 100 μm. n = 3 biological replicates (organoids). ICC staining for ZC4H2 (green).(E) Quantification of pSMAD1/5 protein intensity using a FIJI macro (n = 3 ZARDF organoids). Unpaired t-test. Although a trend toward increased pSMAD1/5 was observed, no significant upregulation of SMAD1/5 phosphorylation was detected in AAV9-treated CtOs compared with UT controls (p value 0.1346).(F) Representative z-stack images of treated AAV9-OE and control GFPKASH-ZARDF CtOs at 4 weeks post-transduction (10x). Scale bar, 100 μm. n = 3 biological replicates. ICC staining for pSMAD1/5 (red).(H) Quantification of MAP2 protein intensity using a FIJI macro (n = 3 ZARDF organoids). Comparison of OE-ZARDF versus UT-ZARDF CtOs by unpaired t-test revealed a significant increase in MAP2 expression (p = 0.0183).(I) Representative z-stack images of treated AAV9-OE and control UT-ZARDF CtOs at 4 weeks post-transduction (10x). Scale bar, 100 μm. n = 3 biological replicates. ICC staining for MAP2 (magenta).(J) Representative z-stack images of treated AAV9-OE and control GFPKASH-ZARDF CtOs at 4 weeks post-transduction taken at 10x. Scale bar, 100 μm. n = 3 biological replicates. ICC staining for MAP2 (magenta).All z-stack images were acquired using an Apotome system and subsequently deconvoluted.

